# A Deterministically Synchronized Widefield Imaging and Virtual Reality Platform for Multimodal Brain–Behavior Recording

**DOI:** 10.64898/2026.05.17.725707

**Authors:** Miguel Maldonado, Omer Faruk Dinc, Macit Emre Lacin, Tenesha Connor, Frederick Bell, Berfin Dinc, Kemal Ozdemirli, Murat Yildirim

## Abstract

**Objective:** Simultaneous recording of brain activity, behaviour, and virtual environments is essential for understanding large-scale neural dynamics during behaviour. However, existing systems often rely on software-based synchronization or post hoc alignment, introducing latency, jitter, and drift that obscure fast brain–behavior interactions.

**Approach:** Here, we present a deterministically synchronized widefield calcium imaging platform that unifies neural imaging, high-speed behavioural monitoring, and closed-loop virtual reality (VR) under a shared hardware-defined clock. This system enables millisecond-precision temporal alignment across modalities, including dual-wavelength hemodynamic correction, pupil and orofacial tracking, locomotion sensing, and VR rendering.

**Main results:** The platform achieves stable hardware-level synchronization across neural imaging, behavioural recordings, and VR rendering without reliance on software timestamps. It supports widefield imaging rates up to 100 Hz and integrates seamlessly with both ViRMEn and Blender VR engines, exhibiting a mean locomotion-to-VR update latency of ∼1.5 ms. Multimodal recordings during VR navigation demonstrate robust temporal alignment between cortical activity, facial dynamics, pupil signals, and locomotion.

**Significance:** This system provides a deterministic multimodal framework for studying brain–behaviour relationships during active behaviour. By enabling millisecond-precision synchronization across neural imaging, behaviour, and virtual environments, this platform enables causal investigation of brain–behaviour interactions at millisecond precision and provides a foundation for next-generation closed-loop neuroengineering experiments.

## 1. Introduction

Large-scale neural dynamics emerge from coordinated activity across distributed cortical networks and evolve on timescales comparable to behavior. Widefield calcium imaging has become a powerful neurophotonics technique for measuring mesoscale brain activity across the dorsal cortex with high spatial coverage and temporal resolution in awake mice^1-3^. By leveraging genetically encoded calcium indicators, widefield imaging enables simultaneous monitoring of activity across multiple cortical regions, providing a systems-level view of neural dynamics underlying sensorimotor processing, cognition, and behavior^1,3^.

When combined with behavioral monitoring and virtual reality (VR), widefield calcium imaging enables controlled investigation of brain-wide activity during locomotion, navigation, and visually guided behavior^4-7^. Head-fixed VR platforms allow precise manipulation of sensory input while maintaining optical stability, making them particularly well suited for optical imaging approaches. Such systems have been widely adopted in studies of spatial navigation, decision-making, and sensorimotor integration using both two-photon and widefield imaging modalities^4-8^.

Accurate interpretation of these experiments critically depends on precise temporal alignment between neural imaging, behavioral measurements, and VR environment updates. Even millisecond-scale timing errors (tens of milliseconds) can distort inferred relationships between cortical activity, motor output, and internal state variables such as pupil-linked arousal and orofacial dynamics^9-12^. As imaging speeds increase and behavioral readouts expand to include multiple high-frame-rate video streams, synchronization errors can obscure fast brain–behavior interactions and introduce ambiguity in causal interpretations.

Existing widefield imaging and VR platforms have enabled important advances in systems neuroscience, yet many rely on software-based synchronization, independent acquisition clocks, or post hoc temporal alignment across modalities^4,6,13^. These approaches can introduce variable latency, frame-to-frame jitter, and timing drift over extended recordings, particularly when multiple cameras, illumination sources, and rendering engines operate concurrently. Moreover, VR environments are often updated through rendering pipelines whose end-to-end latency is rarely quantified, making it difficult to assess the fidelity of closed-loop brain–behavior coupling. Although several systems combine widefield calcium imaging with virtual reality environments, precise temporal synchronization across imaging, behavioural cameras, and VR rendering remains challenging. Small timing mismatches can produce incorrect interpretation of brain–behaviour relationships, particularly for fast cortical dynamics

From a neurophotonics perspective, these limitations are especially problematic for widefield calcium imaging, where accurate separation of calcium-dependent fluorescence from hemodynamic signals and precise alignment with behavior are essential for interpreting fast cortical dynamics^3,14^. Behavioral variables such as locomotion, whisking, and pupil fluctuations are now known to drive widespread cortical activity, often accounting for a substantial fraction of variance in mesoscale signals^9-12^. Disentangling these contributions requires deterministic control over excitation timing, camera exposure, behavioral acquisition, and stimulus updates on a shared temporal reference frame.

Here, we present a fully synchronized widefield calcium imaging platform that integrates brain-wide fluorescence imaging, high-speed behavioral monitoring, and closed-loop virtual reality through deterministic hardware control. The system employs DAQ-driven triggering and microcontroller-based coordination to align widefield imaging with alternating-wavelength excitation for hemodynamic correction, pupil and orofacial videography, locomotion sensing, and VR environment updates. We quantitatively characterize end-to-end timing precision and latency across all subsystems and demonstrate robust performance across multiple imaging rates and VR engines. This platform provides (i) deterministic hardware-level synchronization, (ii) multimodal recordings (brain, pupil, face, locomotion), (iii) compatibility with multiple VR engines, (iv) low-latency closed-loop capability. To our knowledge, this is the first platform to combine deterministic hardware synchronization with multimodal imaging, behavioural tracking, and closed-loop VR while quantitatively validating end-to-end latency and timing precision.

Beyond technical development, we demonstrate the utility of this system by characterizing mesoscale cortical dynamics during behaviour. Specifically, we show that cortical activity is organized into low-dimensional motifs, exhibits indicator-dependent spectral structure, and is strongly predicted by behavioural variables. Together, these results establish deterministic synchronization as a critical requirement for accurately interpreting brain–behaviour relationships and for enabling causal interrogation of brain–behaviour interactions.

## 2. Methods

### 2.1 Overall experimental setup

We designed the experimental platform to achieve deterministic, hardware-level synchronization across widefield calcium imaging, behavioral monitoring, locomotion sensing, and closed-loop virtual reality (VR) rendering (**Fig. 1, Videos 1-2**). The system integrates four primary subsystems: (i) widefield cortical calcium imaging, (ii) infrared video imaging of pupil and orofacial dynamics, (iii) locomotion acquisition using an air-suspended spherical treadmill, and (iv) real-time VR environment rendering. All subsystems were coordinated through a centralized data acquisition (DAQ) architecture to ensure precise temporal alignment without reliance on software timestamps or post hoc synchronization.

**Figure 1.**
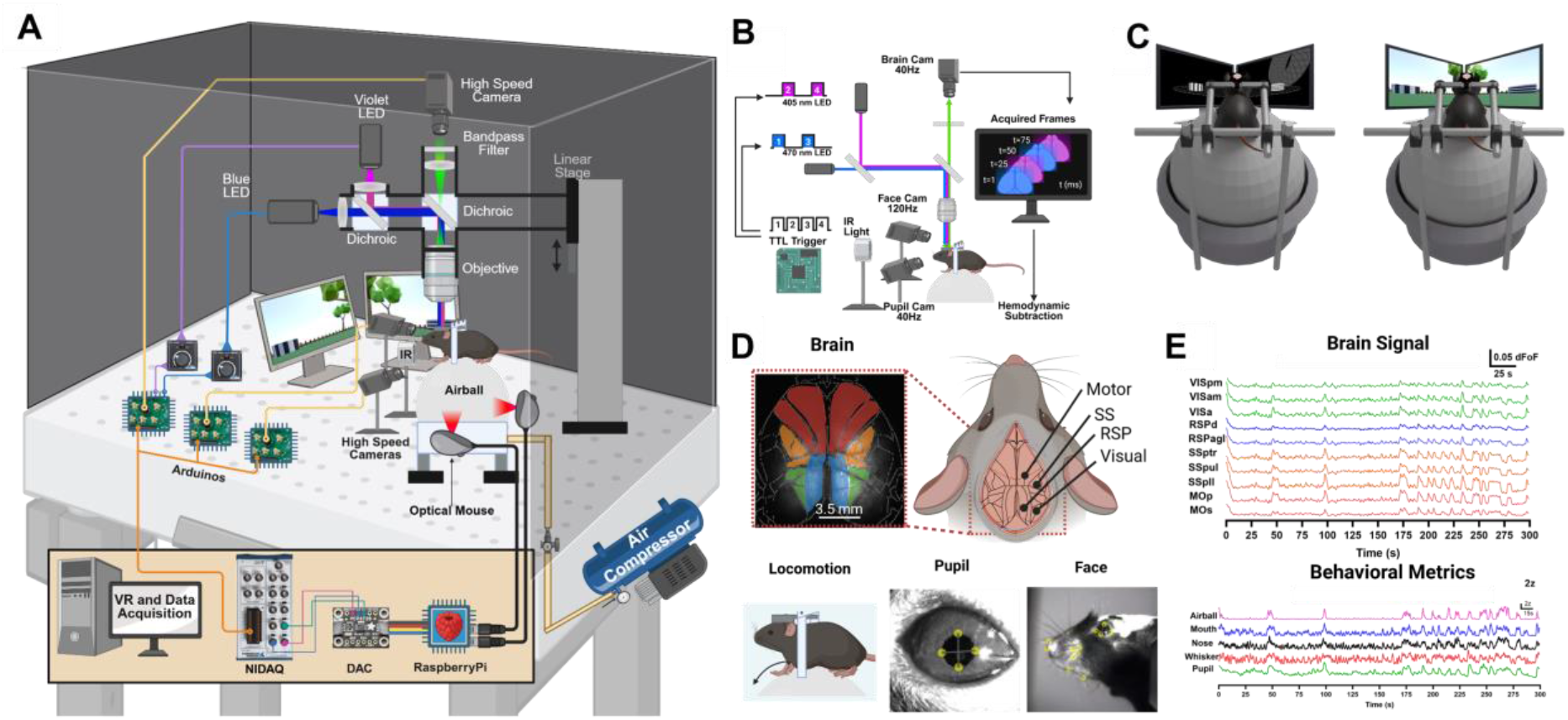
Hardware-synchronized virtual reality and widefield imaging platform for multimodal brain–behavior recording in head-fixed mice. **(A)** Schematic of the integrated experimental system enabling deterministic synchronization across widefield calcium imaging, behavioral monitoring, locomotion sensing, and virtual reality (VR) rendering. The platform combines large-scale one-photon cortical calcium imaging using alternating blue (470 nm) and violet (405 nm) LED excitation, dichroic optics, objective lenses, and a high-speed camera. Camera exposure and LED illumination are controlled via microcontroller-based hardware triggering coordinated by a central data acquisition (DAQ) system. Locomotion is measured using an air-suspended spherical treadmill instrumented with dual USB optical sensors, with signals routed through a Raspberry Pi and DAQ for closed-loop VR control. **(B)** Multicamera configuration for simultaneous acquisition of widefield brain activity (40-100 Hz), pupil dynamics (40 Hz), and orofacial movements (120 Hz) under synchronized fluorescence and infrared illumination. Blue and violet excitation wavelengths are alternated on successive frames to enable separation of calcium-dependent and hemodynamic signals. **(C)** Representative virtual environments used for behavioral experiments, including a minimal high-contrast environment implemented in ViRMEn (left) and a visually complex three-dimensional environment rendered in Blender (right), allowing modulation of visual complexity under identical synchronization constraints. **(D)** Top view of the dorsal cortical surface illustrating anatomical parcellation into visual (VIS), retrosplenial (RSP), somatosensory (SS), and motor (MO) regions, along with example synchronized recordings of locomotion, pupil dynamics, and orofacial features acquired during VR navigation. **(E)** Example widefield calcium activity traces from multiple cortical regions (top) and corresponding behavioral measurements (bottom) recorded simultaneously during VR behavior in a representative wild-type mouse, demonstrating stable temporal alignment across neural and behavioral modalities.

A multifunction DAQ device (PCIe-6353, National Instruments) served as the master timing controller for each experiment. At the start of each recording session, the DAQ generated a digital synchronization signal that initiated acquisition across all subsystems simultaneously. This signal was distributed to multiple microcontroller-based controllers responsible for camera triggering, illumination timing, and behavioral acquisition, ensuring that all data streams shared a common hardware clock throughout the experiment.

We operated widefield calcium imaging, pupil imaging, and orofacial imaging cameras exclusively in external trigger mode, with exposure timing governed by DAQ-generated TTL pulses. We controlled illumination sources for fluorescence excitation and infrared behavioral imaging similarly via hardware triggers to maintain deterministic alignment between excitation, exposure, and acquisition. We acquired locomotion signals from the spherical treadmill digitized by the DAQ and made available in real time to the VR rendering engine, enabling closed-loop updates of the visual environment based on the animal’s movement. An interactive VR landscape rendered from a first-person perspective was projected by two compact liquid crystal display (LCD) projectors (M110, Dell) onto the back of a custom-made 40 cm-diameter translucent acrylic semi-domal screen that was positioned 20 cm in front of the mouse and covered 170° of the mouse’s visual field.

### 2.2 Widefield calcium imaging system

We developed a custom epifluorescence microscope to provide uniform illumination and stable, high-speed imaging across the dorsal cortex for performing widefield calcium imaging. To enable separation of calcium-dependent fluorescence from hemodynamic signals, we delivered excitation light using alternating-wavelength illumination at blue (470 nm; M470L5, Thorlabs) and violet (405 nm; M405LP1, Thorlabs) wavelengths. We combined the two LED sources using a dichroic beamsplitter (87-063, Edmund Optics) and directed onto the cortical surface through a common optical path.

At the sample plane, we maintained average optical power below 10 mW for 470 nm excitation and below 50 mW for 405 nm excitation to minimize photobleaching and phototoxicity while preserving signal-to-noise ratio. We characterized illumination beam profiles using a knife-edge method (1/e^2^ criterion), yielding beam diameters of 9.2 mm (470 nm) and 11.7 mm (405 nm), which provided uniform illumination over an approximately 9.324 × 9.324 mm^2^ field of view encompassing the majority of the adult mouse dorsal cortex.

We collected fluorescence emission through an emission dichroic (T495LPXR, Chroma) and bandpass filter (86-963, Edmund Optics) and imaged onto a monochrome CMOS camera (CS505MU1, Thorlabs) using objective lenses selected to balance field of view, numerical aperture, and acquisition speed. We acquired images at an effective spatial resolution of 252 × 252 pixels with spatial resolution of 0.037 mm/pixel after hardware binning, corresponding to mesoscale cortical coverage suitable for region-level analysis. We performed standard experiments at 40 Hz, with blue and violet excitation alternated on successive frames under deterministic hardware control, ensuring one-to-one correspondence between excitation wavelength and acquired images.

For experiments requiring higher temporal resolution, we enabled an alternative imaging configuration with acquisition rates of up to 100 Hz while preserving full cortical coverage. In this mode, we performed imaging using reduced pixel binning and lower excitation power, without alternating-wavelength excitation, to prioritize temporal resolution over hemodynamic correction.

We governed camera exposure timing and LED switching exclusively by externally generated TTL triggers derived from the master data acquisition (DAQ) clock. This hardware-level control ensured deterministic frame timing, precise alignment between excitation wavelength and image acquisition, and stable synchronization with behavioral and virtual reality subsystems throughout each recording session.

### 2.3 Imaging of pupil and orofacial features

We recorded pupil dynamics and orofacial movements using synchronized infrared (IR) video imaging to enable high-contrast behavioral measurements without interfering with fluorescence excitation. We provided IR illumination by an infrared LED source (LIU780A, Thorlabs) positioned to uniformly illuminate the eye and facial surface.

We acquired pupil images using a monochrome camera (CS505MU1, Thorlabs) equipped with a telecentric lens (58-430, Edmund Optics) to minimize perspective distortion and maintain consistent pupil geometry during small head or eye movements. We captured images at 40 Hz with an effective spatial resolution of 300 × 300 pixels (0.0137 mm/pixel) after 4 × 4 hardware binning, corresponding to a field of view of approximately 4.11 × 4.11 mm^2^. We selected this frame rate and resolution to resolve rapid pupil fluctuations associated with arousal and behavioral state transitions.

We recorded orofacial features, including whisker pad, snout, and jaw movements using a separate monochrome CMOS camera (CS505MU1, Thorlabs) equipped with an infinity-corrected objective (MVL25M23, Navitar). We acquired orofacial images at 120 Hz with an effective resolution of 400 × 400 pixels (0.0588 mm/pixel) after 3 × 3 hardware binning, providing sufficient temporal resolution to capture fast facial movements while maintaining full coverage of the facial region of interest.

We operated both pupil and orofacial cameras exclusively in external trigger mode, with exposure timing governed by DAQ-generated TTL pulses. We synchronized infrared illumination with camera exposure to ensure deterministic temporal alignment with widefield calcium imaging, locomotion sensing, and VR updates.

### 2.4 Locomotion acquisition

We measured animal locomotion using an air-suspended spherical treadmill constructed from lightweight polystyrene and mounted on a custom 3D-printed pedestal. The treadmill allowed unrestricted rotation along forward–backward, lateral, and yaw axes (orbital motion) while maintaining head fixation for optical stability.

We detected ball motion using two orthogonally mounted USB optical motion sensors (Logitech B100, Logitech), which captured translational and rotational components of locomotion. We acquired raw motion data by a Raspberry Pi single-board computer (Raspberry Pi 4 Model B) and converted to analog voltage signals using a digital-to-analog converter (MCP4725, Adafruit). We streamed these analog signals in real time to the data acquisition (DAQ) system.

We digitized locomotion signals by the DAQ and transformed into velocity parameters used to update the animal’s position within the virtual environment. This architecture enabled closed-loop coupling between the animal’s movement and VR rendering while maintaining deterministic timing with respect to neural and behavioral imaging.

### 2.5 Data acquisition and hardware synchronization

A multifunction data acquisition (DAQ) device (PCIe-6353, National Instruments) served as the central timing reference and master clock for all experimental subsystems. The DAQ generated fixed-frequency TTL synchronization pulses (5 V amplitude, 1 ms duration) that initiated and sustained acquisition across widefield imaging, behavioral cameras, locomotion sensing, and virtual reality control.

We distributed synchronization signals from the DAQ to multiple microcontroller-based controllers (Arduino) responsible for routing triggers to cameras and illumination sources. These microcontrollers operated exclusively as deterministic signal routers and did not generate independent clocks, ensuring that all subsystems remained locked to the DAQ timing reference.

We configured all cameras in external trigger mode using manufacturer-provided control software (ThorCam, Thorlabs), with exposure timing defined solely by incoming TTL pulses. We gated LED excitation sources for fluorescence and infrared illumination similarly by hardware triggers to maintain precise alignment between illumination and image acquisition.

This hardware-driven synchronization architecture eliminated reliance on software timestamps or post hoc alignment and minimized latency, jitter, and drift across extended recording sessions. As a result, we recorded neural imaging, behavioral measurements, locomotion signals, and VR environment updates on a shared temporal reference frame, enabling accurate analysis of fast brain–behavior interactions.

### 2.6 Connectivity analysis

We quantified inter- and intra-hemispheric functional connectivity using Pearson correlation applied to Allen atlas–parcellated regional fluorescence time series extracted from the widefield calcium imaging data. For interhemispheric connectivity, we correlated the temporal activity traces of homologous regions in the left and right hemispheres to assess the degree of synchronous activity across hemispheres. In contrast, we calculated intrahemispheric connectivity by computing pairwise Pearson correlations among all regions within the same hemisphere, providing a measure of functional coupling among cortical areas within each hemisphere. We performed connectivity analyses of the hemodynamic-corrected residual signals. We provided the detailed code of this analysis in our github page.

### 2.7 Allen-atlas analysis

To enable atlas-based analysis, we first standardized widefield recordings in orientation using the hemodynamic reference (violet) channel to visually align the cortical midline across animals. Then, we registered the rotated image stacks to the Allen Mouse Brain Common Coordinate Framework (CCFv3) using a landmark-based alignment approach implemented in MATLAB through the locaNMF toolbox. We selected key anatomical landmarks—including anterior cortical anchors near the olfactory bulbs, posterior midline points, and bregma—to guide the alignment. We used these landmarks to compute a similarity transformation that aligned each animal’s dorsal cortical view to the Allen atlas while preserving anatomical geometry. We warped the atlas label map to match the imaging field of view, allowing each pixel to be assigned to a specific cortical parcel. Using these atlas masks, we extracted regional fluorescence time series from the rotated dual-wavelength (470 nm calcium-dependent and 405 nm hemodynamic reference) image stacks by averaging pixel intensities within each region for every frame. Analyses focused on 11 cortical regions including motor (MOp, MOs), retrosplenial (RSPagl, RSPd, RSPv), somatosensory (SSp-ll, SSp-tr, SSp-ul), and visual areas (VISa, VISam, VISpm), with signals computed separately for left and right hemispheres. We provided the detailed code of this analysis in our github page.

### 2.8 Hemodynamic correction

We removed hemodynamic contamination using an atlas-parcellated, region-wise regression approach in which the 405 nm (violet) signal served as a hemodynamic reference for the 470 nm (blue) GCaMP signal. Following alignment and atlas registration, we extracted mean fluorescence time series for each Allen atlas region from both wavelength stacks by averaging pixel intensities within each parcel over time. For each region and hemisphere (left, right, or bilateral), we used the violet trace—dominated by hemodynamic fluctuations related to absorption and scattering—as a regressor to model the hemodynamic component of the blue signal using an ordinary least squares linear regression. We subtracted the predicted hemodynamic contribution from the blue trace, yielding a residual signal representing the hemodynamic-corrected calcium activity for that region. We provided the detailed code of this analysis in our github page.

### 2.9 Motif analysis

We performed motif analysis to identify recurring spatiotemporal patterns of cortical activity using an approach similar to previous mesoscale imaging studies (see e.g., Mohajerani et al., 2020^15^). Prior to analysis, we masked pixels corresponding to the midline and regions outside the brain to restrict the analysis to cortical signals. Because each recording lasted five minutes, we divided the dataset into two equal segments, and we used a 2.5-minute portion for motif extraction. For each pixel, we normalized fluorescence signals and converted to ΔF/F to account for baseline variations across the imaging field. We then analysed the processed imaging data using non-negative matrix factorization (NMF) to decompose the activity into recurring spatiotemporal components representing motifs of cortical activation. We defined motifs as activity patterns with a duration of approximately 2 seconds. For each animal, we quantified the frequency of motif occurrences, and we selected the two most frequently occurring motifs to represent the dominant cortical activation patterns observed during the recording session. We provided the detailed code of this analysis in our github page.

### 2.10 Oscillation analysis

To characterize the spectral composition of the hemodynamic-corrected signals, we quantified oscillatory activity from bilateral cortical region of interest (ROI) time series sampled at 20 Hz. For each region, we estimated the power spectral density using Welch’s method. Then, we integrated spectral power integrated within three frequency bands: delta (0.01–4 Hz), theta (4–8 Hz), and alpha (8–9.99 Hz). To enable comparisons across regions and recordings, we normalized band-specific power values by the total spectral power within the 0.01–9.99 Hz range, yielding relative band power measures. We used these normalized values for all subsequent analyses and visualizations. We provided the detailed code of this analysis in our github page.

### 2.11 Random forest analysis

To quantify the relationship between behavioural variables and neural activity, we applied a supervised machine learning approach based on Random Forest regression. Behavioural predictors included pupil diameter and locomotion speed, which were temporally aligned with the regional brain activity signals. Prior to modelling, we resampled all signals to a common timeline, normalized using z-score normalization, and smoothed using a Savitzky–Golay filter to reduce high-frequency noise. In addition to the raw behavioural variables, we generated several derived features to capture nonlinear and dynamic relationships, including squared terms, interaction terms, rolling variability, and temporal derivatives of pupil and locomotion signals. Then, we trained a Random Forest regression model consisting of multiple decision trees to predict regional brain activity from these behavioural features. We evaluated model performance using out-of-bag prediction and cross-validation to estimate predictive accuracy and generalizability. We assessed feature importance using permutation-based importance measures to determine which behavioural variables contributed most strongly to the prediction of neural activity across brain regions. We provided the detailed code of this analysis in our github page.

### 2.12 Animals

All experiments were conducted in accordance with the guidelines of the Cleveland Clinic Institutional Animal Care and Use Committee (IACUC). Mouse genotyping was performed on ear punches by TransnetYX (Cordova, TN) using their automated real-time PCR platform. Mice were maintained on a 14:10 light: dark cycle with ad libitum access to food and water. The room temperature was regulated between 18 °C and 26 °C. Male and female littermates were separated by sex after weaning and housed with same-sex littermates.

Transgenic reporter mice were generated by crossing either TIT2L-GC6s-ICL-tTA2)-D (stock no. 031562, Jackson Laboratory), or TIGRE2-jGCaMP8m-IRES-tTA2-WPRE (stock no. 037718, Jackson Laboratory) with B6.Cg-Tg(Camk2a-cre)T29-1Stl/J mice (CaMKIIα-Cre, stock no. 005359, Jackson Laboratory).

### 2.13 Surgical Procedures

Cranial window implantation closely followed the procedure previously used for whole-cortex widefield imaging^17^. Mice were anesthetized with isoflurane (induction 2.5%; maintenance 1-1.5%). Buprenorphine (3.25 mg/kg), Meloxicam (5 mg/kg), and sterile saline (0.05 mL/g) were administered at the start of surgery. Anesthesia depth was confirmed by toe pinch. Hair was removed from the dorsal scalp (Nair, Vetiva Mini Hair Trimmer) and the area was disinfected with 3 alternating applications of betadine and 70% isopropanol. Bupivacaine (5 mg/kg) was then injected under the skin for local anesthetic before the scalp was removed. The skull was then exposed, cleaned, and dried. The remaining outer skin was affixed in position with tissue adhesive (Vetbond, 3M) for clean surgical margins. We created an outer wall using dental cement (C&B Metabond, Parkell) while leaving as much of the skull exposed as possible. A custom circular headbar (eMachineShop) was secured in place using dental cement (C&B Metabond, Parkell). A layer of optical glue (Norland Optical Adhesive NOA 81, Norland Products) was then applied to the exposed skull and cured with a UV flashlight (LIGHTFE, UV301Plus-365nm). The mice were allowed to fully recover in a warmed chamber and then returned to their home cages. Post-operational care consisted of three daily injections of Meloxicam (5 mg/kg) following the surgery.

### 2.14 Code and Data availability

Custom hardware designs, microcontroller firmware, acquisition scripts, and analysis codes are freely available in the GitHub repository at https://github.com/yyildirimlab/Widefield-Imaging-and-VR-Platfrom-for-Multimodal-Brain-Behaviour-Recording.git. Datasets generated during this study are available from the corresponding author upon reasonable request.

## 3. Results

### 3.1 Hardware synchronization accuracy across acquisition subsystems

To evaluate temporal alignment across widefield calcium imaging, behavioral monitoring, and virtual reality (VR) control, we directly measured hardware trigger signals and camera exposure timing during synchronized operation. All acquisition subsystems were driven by a shared master clock generated by the data acquisition (DAQ) system, ensuring deterministic timing across modalities (**Fig. 2A**).

**Figure 2.**
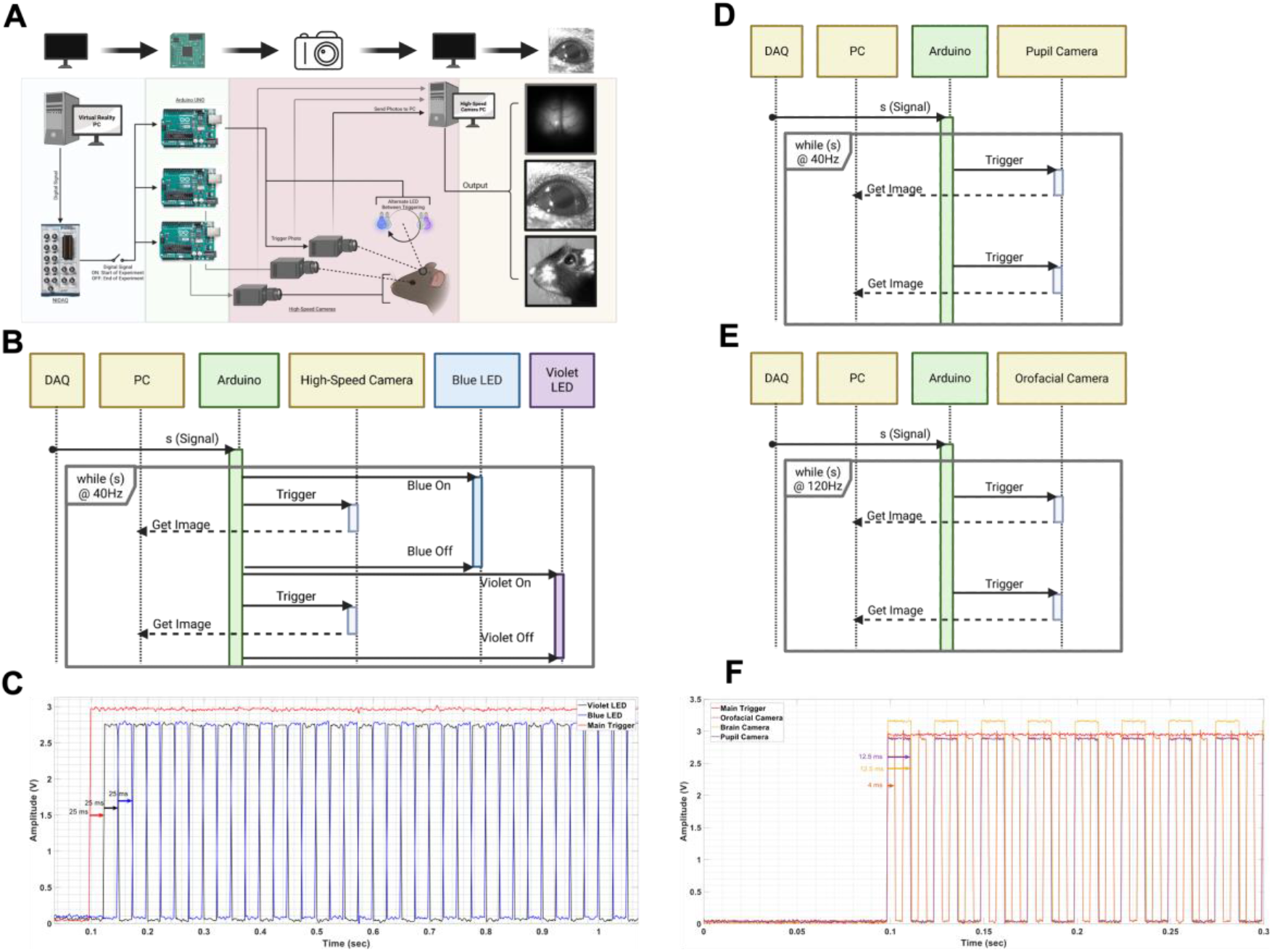
Hardware synchronization and timing relationships across multimodal acquisition subsystems. **(A)** Schematic overview of the synchronized multimodal acquisition architecture integrating widefield calcium imaging, pupil imaging, orofacial imaging, and locomotion sensing during virtual reality (VR). A digital start signal generated by the NI data acquisition (DAQ) system triggers multiple Arduino microcontrollers, which coordinate LED illumination and camera acquisition. High-speed cameras acquire brain, pupil, and facial images under alternating blue (470 nm) and violet (405 nm) excitation to enable separation of calcium-dependent fluorescence from hemodynamic signals. Acquired images are streamed to a dedicated PC for real-time storage. **(B)** Hardware synchronization timeline illustrating the DAQ-driven acquisition loop operating at 40 Hz. Each loop iteration triggers camera exposure and LED illumination in a deterministic sequence, with blue and violet LEDs alternating on successive frames. **(C)** Recorded TTL waveforms showing temporal alignment between the main DAQ trigger and the blue and violet LED control signals. **(D)** Synchronization timeline for the pupil camera operating in external trigger mode, illustrating deterministic alignment with the DAQ trigger. **(E)** Synchronization timeline for the orofacial camera operating at 120 Hz under DAQ-controlled triggering. **(F)** Recorded TTL signals demonstrating simultaneous alignment of brain imaging, pupil camera, and orofacial camera triggers with the main DAQ synchronization signal.

The synchronization architecture operated in a fixed-frequency acquisition loop at 40 Hz, in which each iteration-initiated camera exposure and illumination in a predefined sequence (**Fig. 2B**). We implemented alternating blue (470 nm) and violet (405 nm) excitation on successive frames to enable frame-resolved separation of calcium-dependent fluorescence from hemodynamic signals. Recorded TTL waveforms confirmed precise temporal alignment between the DAQ trigger, LED control signals, and camera exposure, with no drift across acquisition cycles (**Fig. 2C**).

We operated behavioral cameras for pupil and orofacial imaging in external trigger mode and driven by the same DAQ synchronization signal. The pupil camera acquired images at 40 Hz, while the orofacial camera operated at 120 Hz. Timing diagrams and recorded TTL signals demonstrated deterministic alignment between behavioral camera triggers and the master acquisition clock (**Fig. 2D–F**). Across all subsystems, synchronization remained stable throughout experimental sessions, confirming hardware-level temporal alignment without reliance on software timestamps or post hoc correction. Measured trigger timing variability was below the temporal resolution of the acquisition system and did not introduce detectable frame-to-frame jitter.

### 3.2 Characterization of the locomotion-to-virtual-reality control loop and update latency

We next quantified the performance of the closed-loop pipeline linking animal locomotion to VR environment updates. Locomotion signals acquired from the air-suspended spherical treadmill were streamed through the DAQ system and used to drive real-time updates of the visual scene presented to the animal (**Fig. 3A**). Importantly, we initiated VR updates directly from DAQ-available locomotion signals, decoupling VR timing from camera acquisition loops.

**Figure 3.**
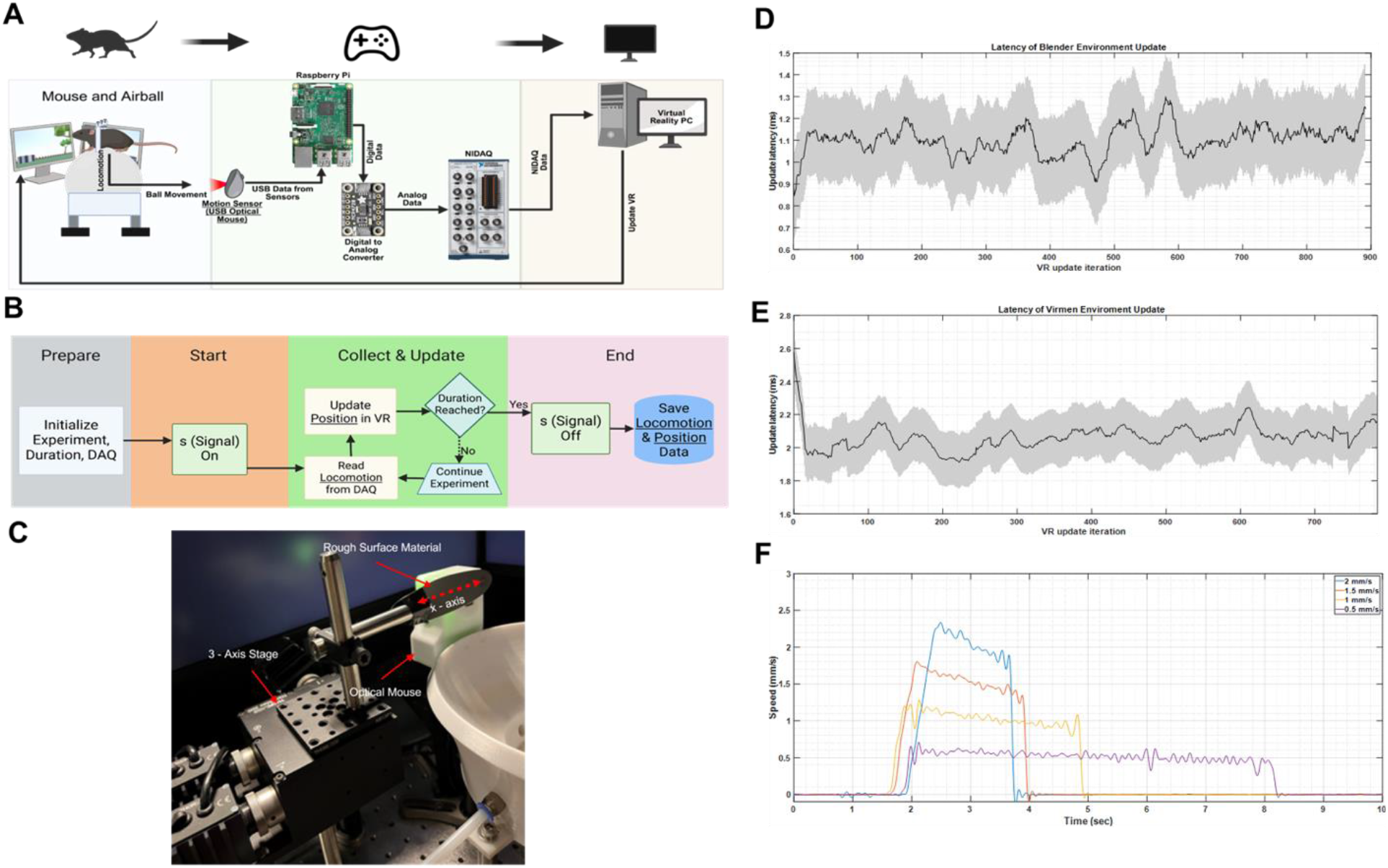
Characterization of the locomotion-to-virtual-reality control loop and update latency. **(A)** Schematic of the closed-loop locomotion-to-virtual-reality (VR) pipeline. Motion of the air-suspended spherical treadmill is detected by optical sensors and transmitted through the data acquisition (DAQ) system to update the virtual environment displayed in front of the animal in real time. **(B)** Algorithmic flow of the VR update loop, illustrating experiment initialization, continuous acquisition of locomotion signals from the DAQ, real-time position updates in the VR environment, and termination upon completion of the experimental duration. **(C)** Characterization of the locomotion sensing system using a three-axis motorized stage (Nanomax 300 Motorized, Thorlabs). A rough-surface material mounted on the stage is translated along the x-axis at controlled speeds to generate reproducible motion inputs for calibration and validation of the optical motion sensors. **(D)** Measured VR update latency during closed-loop operation using the Blender-based VR environment, quantified as the time delay between locomotion signal availability at the DAQ input and issuance of the corresponding VR update command. **(E)** Measured VR update latency using the ViRMEn-based VR environment, demonstrating comparable latency profiles under identical measurement conditions. **(F)** Resulting locomotion speed profiles obtained from the calibration procedure in (C), showing responses at different translational speeds over the same travel distance, confirming consistent sensor behavior across operating regimes.

The VR control pipeline followed a deterministic processing sequence consisting of experiment initialization, continuous acquisition of locomotion signals, real-time position updates within the virtual environment, and termination upon completion of the predefined recording duration (**Fig. 3B**). This architecture ensured consistent handling of locomotion inputs throughout each experimental session.

To validate locomotion sensing accuracy and characterize system response, we calibrated optical motion sensors using a three-axis motorized translation stage. We translated a rough-surface material mounted on the stage along the x-axis at controlled velocities to generate reproducible motion inputs (**Fig. 3C**). Across tested speeds, the resulting velocity profiles were stable and reproducible, confirming reliable sensor performance and linear response over the operating range (**Fig. 3F**).

We quantified end-to-end VR update latency during closed-loop operation by measuring the time delay between locomotion signal availability at the DAQ input and issuance of the corresponding VR update command. Using a Blender-based VR environment, the system exhibited a mean update latency of approximately 1.5 ms with minimal variability across successive updates (**Fig. 3D**). We observed comparable latency distributions when using a ViRMEn-based VR environment under identical conditions (**Fig. 3E**), demonstrating that the hardware-driven control architecture provides VR engine–agnostic timing behavior.

### 3.3 Demonstration of synchronized neural and behavioural acquisition during virtual reality navigation

To demonstrate system performance during representative experiments, we acquired synchronized neural and behavioral data during head-fixed navigation in visually distinct VR environments. Experimental workflows included developing transgenic mice with genetically encoded calcium indicators, headplate implantation, and subsequent navigation in VR environments rendered using either ViRMEn or Blender (**Fig. 4A, D**).

**Figure 4.**
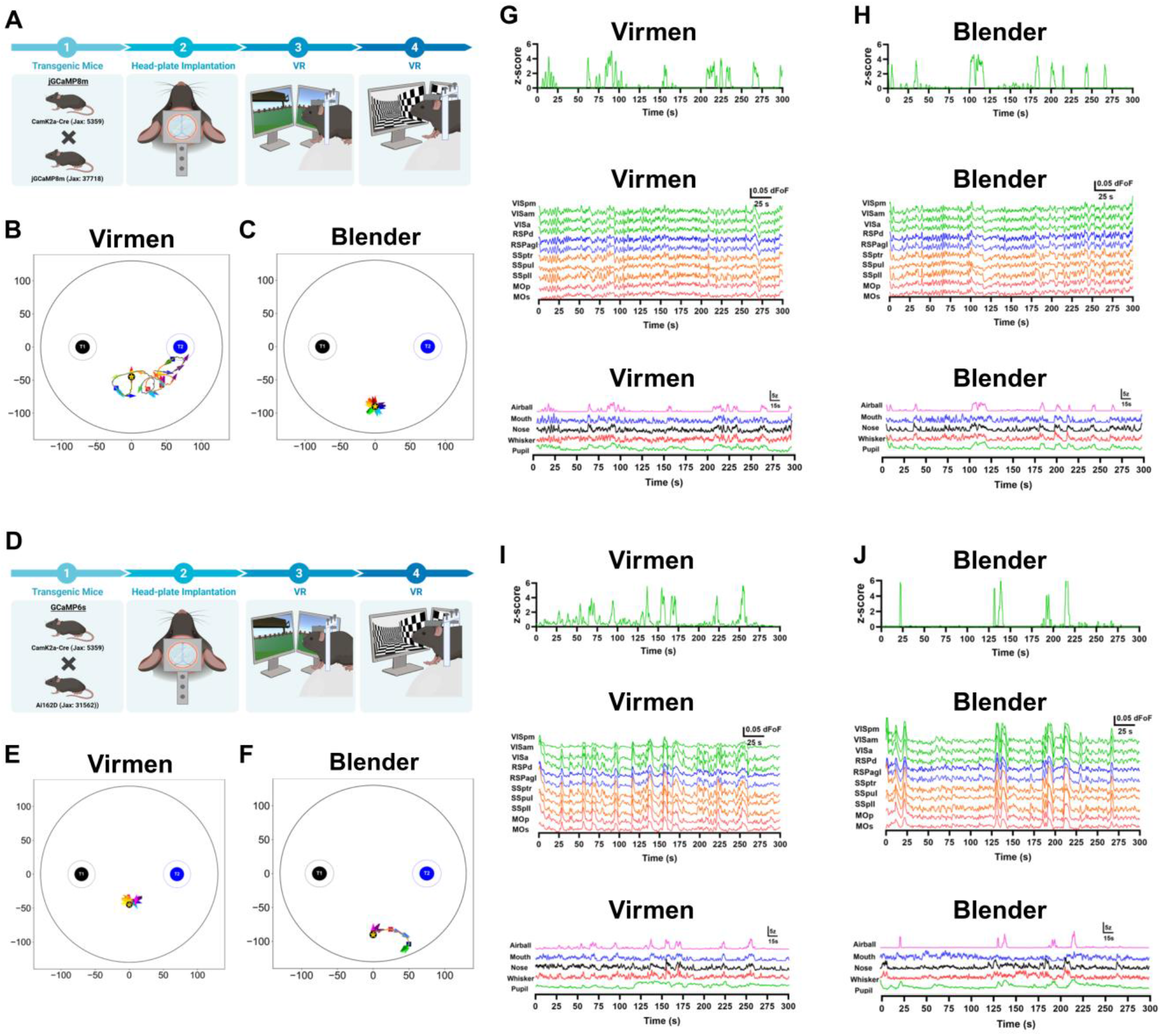
Brain-wide neural dynamics and behavior across visually distinct virtual reality environments. (**A**,**D**) Experimental workflow illustrating GCaMP6s and jGCaMP8m, headplate implantation, and subsequent head-fixed navigation in virtual reality (VR) environments rendered using ViRMEn (high-contrast black-and-white) and Blender (realistic, colorful). (**B-C, E-F**) Example locomotion trajectories and corresponding instantaneous locomotion speeds acquired from two optical motion sensors during navigation in the ViRMEn (B,E), and in the Blender (C,F) environments, respectively. (**G–H, I-J**) Representative synchronized recordings from the same animal during navigation in ViRMEn (G,I) and Blender (H,J). Traces show locomotion signals (top), widefield calcium activity from multiple cortical regions (middle), and behavioral metrics including pupil dynamics and orofacial motion (bottom), demonstrating stable temporal alignment across neural and behavioral modalities under distinct VR visual conditions.

Locomotion behavior recorded using the spherical treadmill exhibited stable trajectories and consistent speed profiles across both VR environments (**Fig. 4B-C, E-F**), indicating reliable closed-loop control independent of visual scene complexity. Neural and behavioral data were acquired simultaneously under identical synchronization constraints in both environments.

Representative recordings from the same animal revealed stable temporal alignment between locomotion, widefield calcium activity across multiple cortical regions, pupil dynamics, and orofacial movements during navigation in both ViRMEn and Blender environments (**Fig. 4G-J**). Neural signals remained precisely aligned with behavioral measurements throughout the recording sessions, demonstrating that deterministic synchronization was preserved during extended closed-loop experiments.

Together, these results confirm that our platform supports robust, synchronized acquisition of brain-wide neural activity and high-speed behavioral dynamics during VR-guided behavior. These findings demonstrate that deterministic synchronization is essential for preserving temporal fidelity in multimodal recordings, enabling accurate interpretation of fast brain–behavior interactions that would otherwise be obscured by timing variability. The consistency of temporal alignment across modalities and VR engines establishes the system as a reliable infrastructure for investigating mesoscale brain–behavior coupling under controlled sensory conditions. This level of temporal fidelity is particularly important for resolving fast brain–behavior interactions, including arousal- and movement-related cortical dynamics, where millisecond-scale timing errors can obscure causal relationships.

### 3.4 Interhemispheric and intrahemispheric cortical connectivity across imaging indicators

To characterize large-scale cortical communication across hemispheres and within individual hemispheres, we quantified functional connectivity between predefined cortical regions during behaviour. We first examined interhemispheric connectivity by calculating the coefficient of determination (R^2^) between homologous cortical regions across the left and right hemispheres, including motor (MO), primary somatosensory (SSp), retrosplenial (RSp), and visual (VIS) cortices (**Fig. 5A–B**). We performed this analysis independently in mice expressing GCaMP6s (**Fig. 5C**) and jGCaMP8m (**Fig. 5D**) to determine whether connectivity patterns were consistent across calcium indicators.

**Figure 5.**
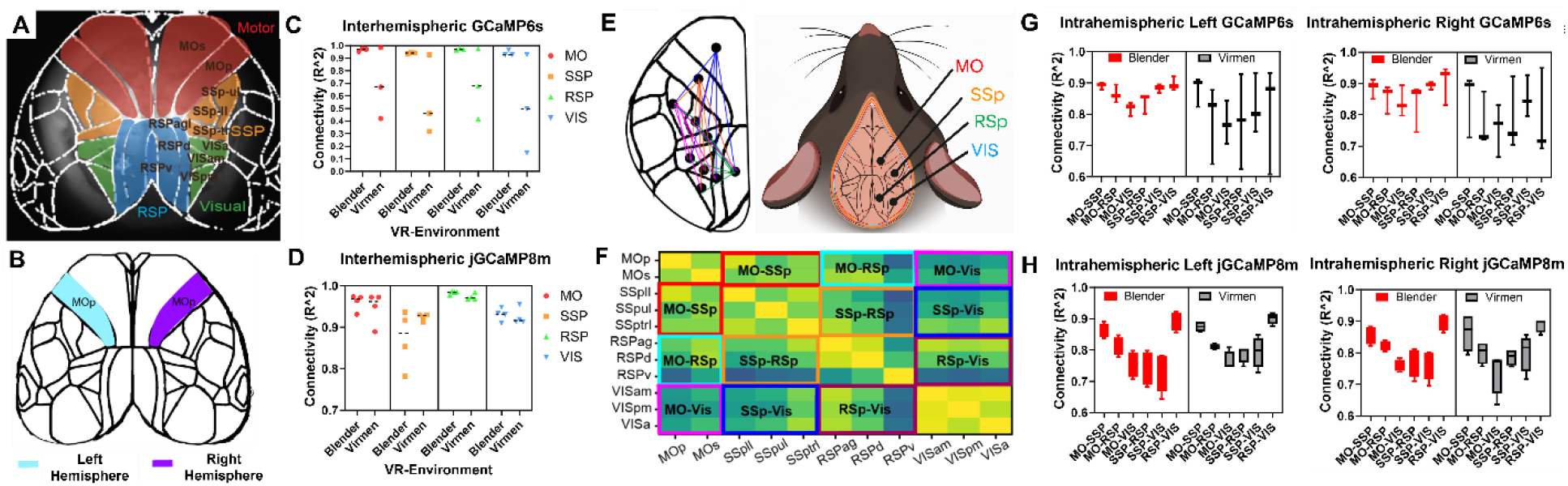
Interhemispheric and intrahemispheric functional connectivity across cortical regions in GCaMP6s and jGCaMP8m mice. **(A)** Schematic representation of the cortical regions used for connectivity analysis, including motor (MO), primary somatosensory (SSp), retrosplenial (RSp), and visual (VIS) areas mapped onto the dorsal cortical surface. **(B)** Example anatomical segmentation illustrating homologous regions in the left and right hemispheres, shown here for the primary motor cortex (MO). **(C)** Interhemispheric connectivity (R^2^) between homologous cortical regions in GCaMP6s mice (n=3) during different virtual reality (VR) environments. **(D)** Interhemispheric connectivity analysis for the same cortical regions in jGCaMP8m mice (n=4). **(E)** Schematic representation of the intrahemispheric connectivity analysis. Left: individual cortical regions used for pairwise connectivity calculations. Right: grouping of cortical regions into four functional clusters (MO, SSp, RSp, VIS). **(F)** Matrix representation illustrating how pairwise intrahemispheric connectivity values were grouped to compute connectivity within and between the four cortical clusters. **(G)** Intrahemispheric connectivity results for GCaMP6s mice (n=3), shown separately for the left hemisphere (left) and right hemisphere (right). **(H)** Intrahemispheric connectivity results for jGCaMP8m mice (n=4), shown separately for the left hemisphere (left) and right hemisphere (right). In G-H, red dots present Blender and black dots present Virmen environments.

Next, we evaluated intrahemispheric connectivity within each hemisphere. We grouped individual cortical regions into four functional clusters (MO, SSp, RSp, and VIS) and computed pairwise correlations between regions within the same hemisphere (**Fig. 5E–F**). Then, we summarized connectivity values for each hemisphere to assess network organization within the left and right cortical hemispheres. These analyses revealed consistent intrahemispheric connectivity patterns across animals for both GCaMP6s (**Fig. 5G, Videos 3-4**) and jGCaMP8m mice (**Fig. 5H, Videos 5-6**), demonstrating that large-scale cortical interactions are robust across imaging indicators and hemispheric networks.

### 3.5 Mesoscale cortical dynamics are organized into recurring spatiotemporal motifs across environments and calcium indicators

To characterize recurring large-scale cortical activity patterns during behaviour, we applied a motif extraction approach to mesoscale calcium imaging recordings (**Fig. 6A**). This analysis decomposes neural activity into spatial motifs and their associated temporal weight profiles, enabling identification of dominant spatiotemporal activity patterns that recur throughout the recording. Using a deconvolution-based approach, we extracted motifs that capture the principal modes of cortical dynamics while preserving both spatial organization and temporal contributions of each motif^15^.

**Figure 6.**
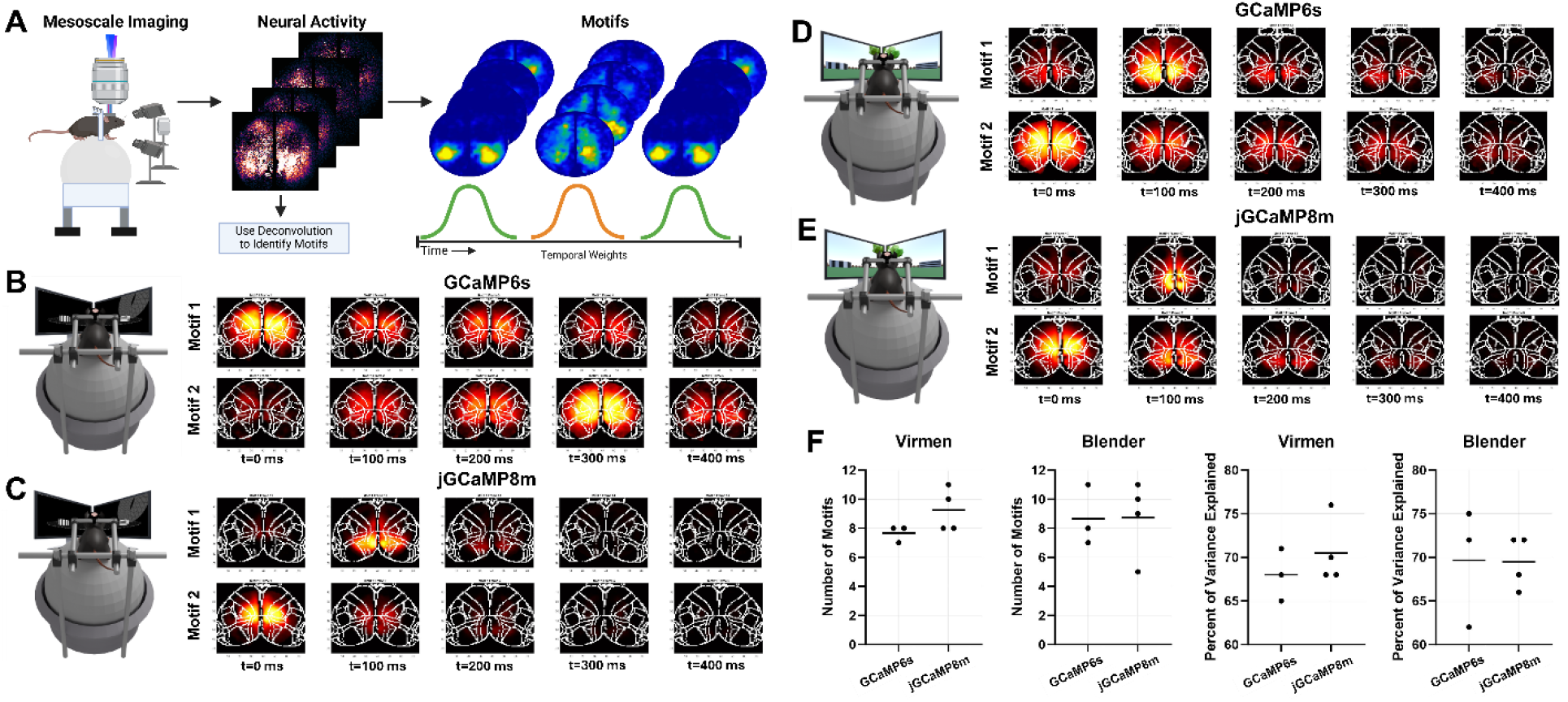
Spatiotemporal cortical activity motifs extracted from mesoscale calcium imaging across different indicators and behavioral environments. **(A)** Schematic overview of the motif extraction pipeline. Mesoscale calcium imaging recordings were acquired during behavior, generating large-scale cortical activity maps over time. Deconvolution-based analysis was used to identify recurring spatiotemporal activity motifs, consisting of spatial activation patterns and associated temporal weight profiles that describe their occurrence across time using nonnegative matrix factorization^14^. **(B)** Representative motifs extracted from GCaMP6s mice during navigation in the ViRMEn virtual reality environment, showing the two most frequently occurring motifs across animals. **(C)** Representative motifs extracted from jGCaMP8m mice in the ViRMEn environment, illustrating the two dominant motifs observed across recordings. **(D)** Representative motifs extracted from GCaMP6s mice during behavior in the Blender-based virtual environment, highlighting the two most prevalent motifs. **(E)** Representative motifs extracted from jGCaMP8m mice in the Blender environment, again showing the two dominant motifs identified from the data. **(F)** Summary statistics of motif analysis across imaging indicators and environments. Left panels show the number of motifs identified in ViRMEn and Blender environments. Right panels show the percentage of variance explained by the extracted motifs for both GCaMP6s and jGCaMP8m datasets.

We first examined motif structure during navigation in the ViRMEn virtual reality environment. In GCaMP6s mice, the two most frequently occurring motifs revealed distinct patterns of cortical engagement (**Fig. 6B**). One motif predominantly activated motor cortical regions, whereas the second motif involved a broader network encompassing motor, primary somatosensory, and retrosplenial cortices, suggesting coordinated activity across sensorimotor and associative areas during navigation. In contrast, jGCaMP8m mice exhibited motifs with slightly different spatial profiles (**Fig. 6C**). One motif showed dominant activation in the retrosplenial cortex, while the second motif primarily engaged motor regions, indicating that navigation-related cortical dynamics in these animals were largely organized around motor and retrosplenial activity patterns.

We next examined motif organization during behaviour in the Blender-based virtual environment. In GCaMP6s mice, the dominant motifs consisted of one motif primarily activating the retrosplenial cortex and another motif engaging motor regions (**Fig. 6D**). A similar motif structure was observed in jGCaMP8m mice, where one motif was characterized by strong retrosplenial activation, while the second motif primarily involved motor cortical regions (**Fig. 6E**). These results suggest that across environments and calcium indicators, cortical activity motifs are largely organized around recurring activation of motor and retrosplenial networks, with occasional engagement of additional sensorimotor regions.

To quantify motif structure across conditions, we compared the number of motifs identified and the percentage of variance explained by these motifs across environments and calcium indicators (**Fig. 6F**). The number of extracted motifs was similar between GCaMP6s and jGCaMP8m mice in both environments, indicating comparable motif complexity across indicators. Moreover, a relatively small number of motifs accounted for a substantial proportion of the variance in cortical activity, demonstrating that mesoscale cortical dynamics during behaviour can be effectively summarized by a limited set of recurring spatiotemporal patterns. This suggests that large-scale cortical activity during behaviour is organized into a low-dimensional set of recurring network states, rather than independent activity across regions, highlighting the dominant role of motor and retrosplenial circuits in coordinating navigation-related dynamics.

### 3.6 Spectral analysis reveals indicator-dependent differences in cortical dynamics

To investigate the spectral characteristics of large-scale cortical dynamics, we analysed mesoscale calcium signals from defined cortical regions using frequency-domain analysis. Example cortical regions were selected based on the Allen Brain Atlas, including the secondary motor cortex (MOs) and visual cortex (VISa) (**Fig. 7A**). Representative activity traces from these regions revealed fluctuations in calcium signals during behaviour, reflecting ongoing mesoscale neural dynamics. To quantify the frequency content of these signals, we applied Fourier transform analysis to convert the time-domain activity traces into their corresponding frequency spectra (**Fig. 7B**).

**Figure 7.**
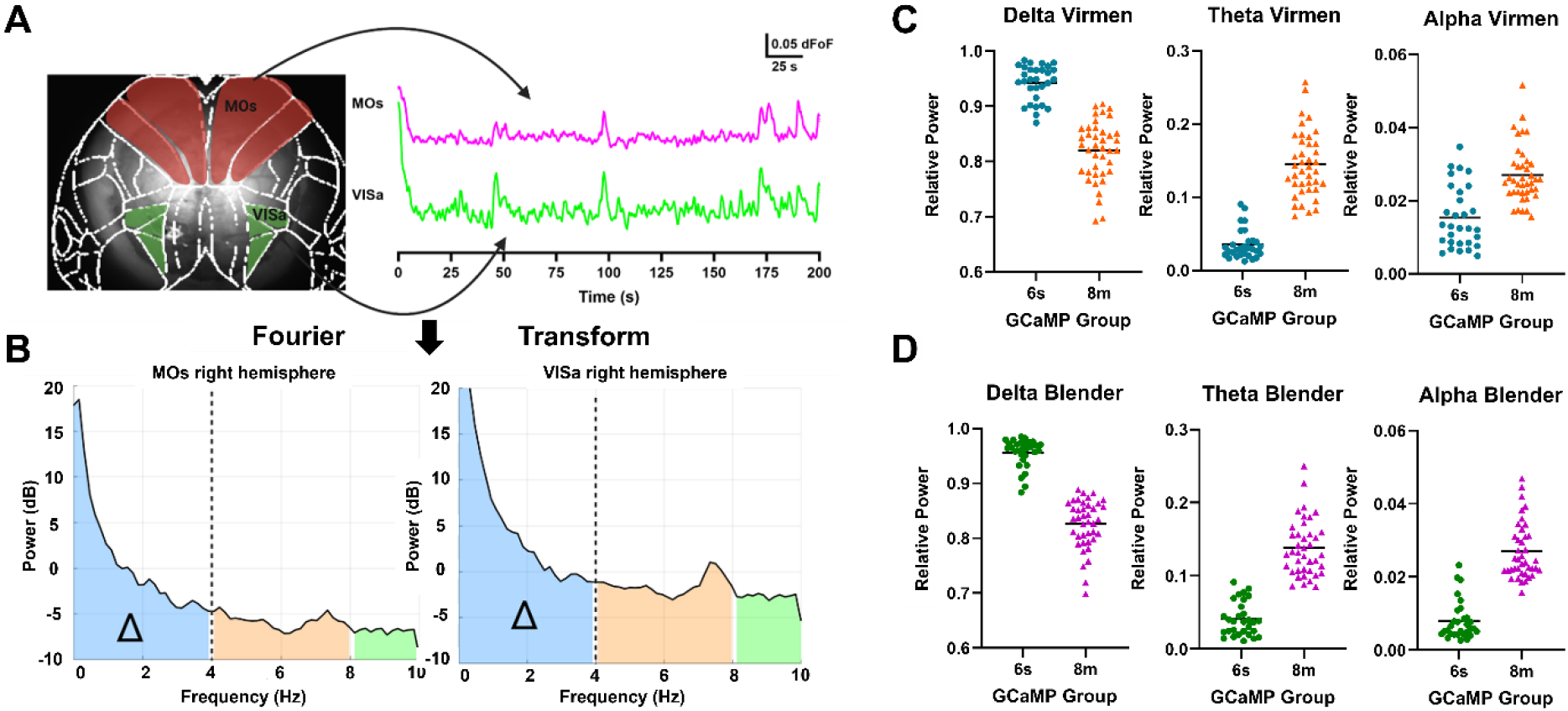
Frequency-domain analysis of cortical activity across calcium indicators and behavioral environments. **(A)** Example cortical regions and activity traces used for spectral analysis. Left: Allen Brain Atlas reference showing the secondary motor cortex (MOs) and visual cortex (VISa). Right: representative fluorescence activity traces (ΔF/F) from MOs and VISa regions during behavior. **(B)** Fourier transform analysis of cortical activity signals from MOs and VISa regions, illustrating the power spectrum of neural activity across frequencies. Shaded regions indicate frequency bands corresponding to delta (0–4 Hz), theta (4–8 Hz), and alpha (8–10 Hz). **(C)** Relative power of delta, theta, and alpha frequency bands in the ViRMEn virtual environment for GCaMP6s and jGCaMP8m mice. Each point represents an individual brain activity recording session. **(D)** Relative power of delta, theta, and alpha frequency bands in the Blender virtual environment for GCaMP6s and jGCaMP8m mice. Across both environments, jGCaMP8m mice exhibit reduced relative delta power and increased power in higher frequency bands compared with GCaMP6s mice.

We next quantified the relative power within canonical frequency bands, including delta (0–4 Hz), theta (4–8 Hz), and alpha (8–10 Hz), across experimental conditions. In the ViRMEn virtual environment, we observed consistent differences between calcium indicators (**Fig. 7C**). Specifically, jGCaMP8m mice exhibited lower relative power in the delta frequency band compared with GCaMP6s mice, while showing higher relative power in theta and alpha frequency ranges. A similar trend was observed in the Blender environment (**Fig. 7D**), where jGCaMP8m mice again showed reduced delta power and increased higher-frequency power relative to GCaMP6s mice. These results indicate that spectral properties of mesoscale cortical activity vary across calcium indicators, with jGCaMP8m signals emphasizing higher-frequency components of cortical dynamics, while GCaMP6s recordings exhibit relatively stronger low-frequency activity. These results indicate that calcium indicators differentially capture temporal features of cortical activity, suggesting that indicator kinetics influence the observed frequency structure of mesoscale dynamics and should be considered when interpreting or comparing datasets.

### 3.7 Behavioural variables predict mesoscale cortical activity across brain regions

To quantify the relationship between behavioural dynamics and mesoscale cortical activity, we trained random forest regression models to predict neural activity from behavioural variables. Behavioural inputs included pupil diameter as well as locomotion variables derived from the spherical treadmill (**Fig. 8A**). We used these behavioural features as predictors to estimate cortical activity for individual brain regions. We quantified the model performance using the coefficient of determination (R^2^) between predicted and ground-truth calcium signals.

**Figure 8.**
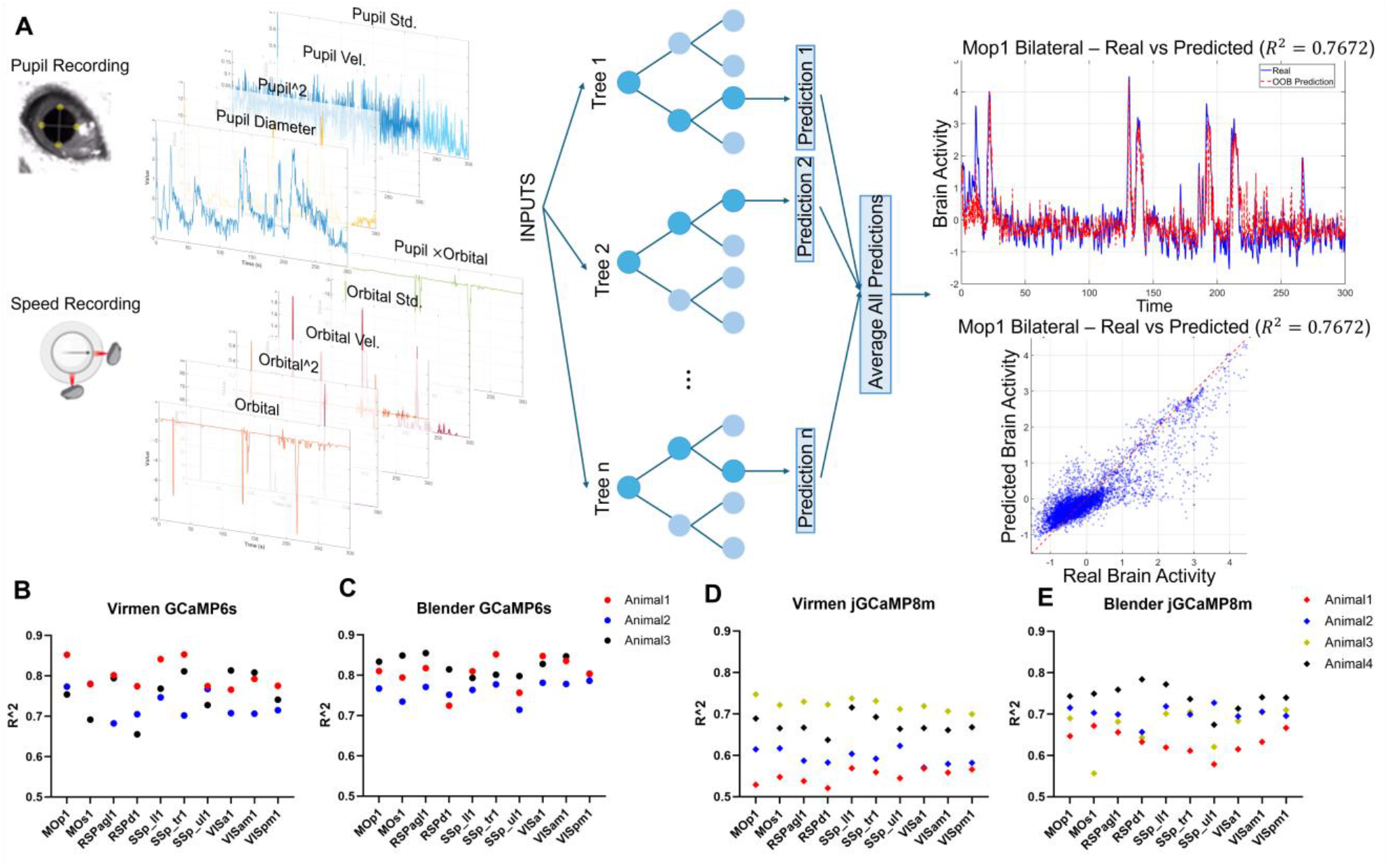
Prediction of mesoscale cortical activity from behavioral variables using random forest regression. **(A)** Schematic of the behavioral-to-brain prediction framework. Pupil diameter and locomotion variables derived from the spherical treadmill (including orbital speed) were extracted from behavioral recordings and used as inputs to a random forest regression model to predict cortical activity. Model performance was evaluated by comparing predicted neural activity with ground-truth calcium signals from individual cortical regions. **(B)** Prediction performance for GCaMP6s mice in the ViRMEn virtual reality environment, shown as coefficient of determination (R^2^) values across cortical regions. Results are shown separately for each animal for bilateral hemisphere predictions. Each point represents the prediction performance for an individual animal. **(C)** Prediction performance for GCaMP6s mice in the Blender environment, with R^2^ values shown across cortical regions for bilateral hemisphere predictions. **(D)** Prediction performance for jGCaMP8m mice in the ViRMEn environment, showing model accuracy across animals for bilateral hemisphere. **(E)** Prediction performance for jGCaMP8m mice in the Blender environment, again showing prediction accuracy across animals for bilateral hemisphere activity.

We first evaluated prediction performance in the ViRMEn virtual reality environment. In GCaMP6s mice, behavioural variables predicted bilateral (average of left and right hemisphere) cortical activity with relatively high accuracy across multiple cortical regions, with R^2^ values typically ranging between approximately 0.65 and 0.80 (**Fig. 8B, Video 7**). In jGCaMP8m mice, the random forest models also predicted bilateral cortical activity across regions, although the R^2^ values were generally slightly lower compared with GCaMP6s recordings (**Fig. 8D, Video 8**).

We next examined prediction performance in the Blender virtual environment. In GCaMP6s mice, behavioural variables again predicted bilateral cortical activity across brain regions with similar accuracy to that observed in the ViRMEn environment (**Fig. 8C, Video 9**). In jGCaMP8m mice, prediction performance remained robust across cortical areas and hemispheres (**Fig. 8E, Video 10**), though R^2^ values were again modestly lower than those observed for GCaMP6s recordings. Together, these results demonstrate that behavioural variables such as pupil dynamics and locomotion strongly predict large-scale cortical activity, highlighting the tight coupling between behavioural state and mesoscale neural dynamics across experimental environments. Together, these findings support a framework in which behavioural and internal state variables account for a substantial fraction of cortex-wide activity, emphasizing the importance of precise temporal alignment for accurately quantifying brain–behaviour relationships.

## 4. Discussion

This study presents a deterministically synchronized widefield calcium imaging and virtual reality (VR) platform that enables precise alignment of brain-wide neural activity with behavior in head-fixed mice. To our knowledge, this is the first system to unify neural imaging, dual-wavelength hemodynamic correction, high-speed behavioral monitoring, and closed-loop VR under a shared hardware-defined clock with quantitative validation of timing performance. Recent work has shown that a substantial fraction of cortex-wide activity can be explained by spontaneous movements and arousal fluctuations rather than task structure alone^3,9,16^. By synchronizing widefield imaging with high-speed recordings of locomotion, pupil dynamics, and orofacial movements, our platform enables rigorous separation of these contributions and supports more accurate interpretation of mesoscale brain dynamics.

A key advantage of this architecture is its ability to support multimodal analyses of mesoscale cortical dynamics during behavior. Using the synchronized dataset generated by the platform, we characterized large-scale cortical activity at multiple levels of organization. Functional connectivity analysis revealed stable interhemispheric and intrahemispheric interactions across motor, somatosensory, retrosplenial, and visual regions, indicating robust coordination of distributed cortical networks during virtual navigation. At the level of population dynamics, motif analysis identified recurring spatiotemporal activation patterns dominated by motor and retrosplenial cortical engagement, consistent with the roles of these regions in locomotion and spatial navigation. Spectral analysis further revealed differences in frequency structure between calcium indicators, with jGCaMP8m recordings emphasizing higher-frequency components of cortical activity relative to GCaMP6s signals. Finally, machine learning models demonstrated that behavioral variables such as pupil fluctuations and locomotion speed predict a substantial portion of mesoscale cortical activity across brain regions. Together, these analyses illustrate how synchronized multimodal recordings enable integrated characterization of connectivity, dynamics, and behavioral coupling within large-scale cortical networks.

In particular, pupil-linked arousal and locomotion exert rapid and dissociable effects on cortical activity and sensory processing^**10-12**^. The deterministic timing achieved here is essential for resolving such fast interactions, as even small timing errors can obscure causal relationships between behavior and neural activity. The ability to combine dual-wavelength excitation for hemodynamic correction with high-speed behavioral monitoring further strengthens the biological interpretability of widefield signals^**17**^.

From an engineering standpoint, the principal contribution of this work is a hardware-driven synchronization architecture in which all acquisition and control subsystems operate on a shared DAQ-defined clock. Unlike software-based or post hoc synchronization approaches commonly used in widefield and VR experiments^4,18^, our system minimizes latency, jitter, and drift across extended recordings. We explicitly quantified the closed-loop locomotion-to-VR update latency (∼1.5 ms) and demonstrated stable timing across two distinct VR engines (ViRMEn and Blender), highlighting the engine-agnostic nature of the platform. Quantitative characterization of timing performance remains rare in behavioral imaging systems and represents an important step toward reproducible neurophotonics instrumentation. By eliminating reliance on software timestamps and post hoc alignment, this approach establishes deterministic synchronization as a key design principle for next-generation multimodal neuroimaging systems.

Relative to existing VR-based imaging platforms^4-6,8^, our system uniquely integrates widefield calcium imaging, dual-wavelength hemodynamic correction, multi-camera behavioral monitoring, and VR control under deterministic hardware synchronization. While prior platforms have enabled landmark discoveries in navigation and sensorimotor integration^4-6^, they often rely on independent clocks or unreported display latencies. By unifying all data streams on a single temporal reference, the present platform improves fidelity and reproducibility for experiments probing fast brain–behavior coupling (Table 1).

**Table 1.** Comparison of experimental platforms for neural recording during behaviour and virtual environments.

| System / Paper | Neural recording | Cortical Imaging Area (mm<sup>2</sup>) | Orofacial Tracking | VR Display FOV (deg) | Neural Recording Rate(Hz) | Latency (ms) | Notes / Distinct Features |
| --- | --- | --- | --- | --- | --- | --- | --- |
| **Maldonado et al. (2026)** | 1-Photon Widefield calcium imaging | 9.324 × 9.324 ≈ 87 mm<sup>2</sup> (full dorsal cortex) | Yes | - | 40–100 Hz (dual LED 470 + 405 nm) | ~1.5 ms | Synchronized brain, pupil (40 Hz), face (120 Hz), locomotion; multimodal deep learning analysis. |
| **Hayashi et al. (2017)** | 1-Photon with GRIN LENS | <1 mm<sup>2</sup> | No | 90° h × 77° v (flat panel array) | 10 Hz | Not Reported | GRIN lens-based imaging enabling access to deep brain structures. |
| **Lopes et. Al. (2021)** | Electrophysiology | Not applicable | No | Display – dependent | 60 Hz | ~20-40 ms | Flexible VR framework: electrophysiology-focused; no optical imaging or face tracking |
| **West et al. (2022)** | 1-Photon Widefield calcium imaging | 6.2 × 6.2 ≈ 38.4 mm<sup>2</sup> | No | No | 20 - 40 Hz | Not applicable | Mesoscopic cortical imaging during locomotion; no closed-loop VR environment |

The system also provides a strong foundation for future closed-loop extensions, including real-time modulation of VR environments or optogenetic perturbations triggered by neural or behavioral states. Low-latency feedback has proven critical for causal interrogation of neural circuits^19^, and the hardware architecture described here is well suited for such applications. More broadly, this platform enables closed-loop experimental paradigms in which neural or behavioral signals can dynamically drive sensory input or targeted perturbations, providing a foundation for causal interrogation of large-scale neural circuits. Such capabilities are critical for advancing both basic neuroscience and translational applications aimed at restoring brain function in neurological disorders^20-23^.

## 5. Limitations

Several limitations should be considered when interpreting these results. First, the platform relies on a head-fixed preparation, which restricts the range of natural behaviors and may influence internal state. While head fixation is essential for stable widefield imaging, findings may not fully generalize to freely moving conditions. Second, the use of alternating-wavelength excitation for hemodynamic correction imposes a tradeoff between temporal resolution and signal fidelity, effectively reducing the sampling rate of calcium-dependent fluorescence. Although sufficient for mesoscale dynamics, this approach may limit sensitivity to very fast neural events. Finally, the system represents a custom-built, single-animal setup, and broader adoption will require technical expertise and potential adaptation to other species, behaviors, or sensory modalities.

## 6. Conclusion

In summary, we developed a deterministically synchronized widefield calcium imaging and VR platform that enables precise, reproducible measurement of mesoscale brain activity during behaviour. By unifying all subsystems under a shared timing architecture, this platform overcomes key limitations of existing approaches and enables accurate investigation of fast brain–behaviour interactions. This work provides a scalable framework for multimodal neurophotonics and lays the groundwork for closed-loop and disease-focused studies aimed at causally linking large-scale neural dynamics to behaviour and enabling targeted interventions in health and disease.

## Acknowledgements

This work was supported by US National Institute of Health (NIH) grants #R00EB027706 (MY), Case Comprehensive JumpStart Grant #3209 (MY), American Brain Tumor Association Discovery Grant #DG2500080 in memory of Kaitlyn Berg (MY), Cleveland Clinic Research Startup (MY), Case Western University SOURCE Fellowship (KO, FB).

## References

[1] Ren, C. & Komiyama, T. Characterizing Cortex-Wide Dynamics with Wide-Field Calcium Imaging. J Neurosci 41, 4160–4168 (2021). 10.1523/JNEUROSCI.3003-20.2021

[2] Nietz, A. K. et al. Wide-Field Calcium Imaging of Neuronal Network Dynamics In Vivo. Biology (Basel) 11 (2022). 10.3390/biology11111601

[3] Musall, S., Kaufman, M. T., Juavinett, A. L., Gluf, S. & Churchland, A. K. Single-trial neural dynamics are dominated by richly varied movements. Nat Neurosci 22, 1677–1686 (2019). 10.1038/s41593-019-0502-4

[4] Aronov, D. & Tank, D. W. Engagement of neural circuits underlying 2D spatial navigation in a rodent virtual reality system. Neuron 84, 442–456 (2014). 10.1016/j.neuron.2014.08.042

[5] Dombeck, D. A., Harvey, C. D., Tian, L., Looger, L. L. & Tank, D. W. Functional imaging of hippocampal place cells at cellular resolution during virtual navigation. Nat Neurosci 13, 1433–1440 (2010).

[6] Harvey, C. D., Coen, P. & Tank, D. W. Choice-specific sequences in parietal cortex during a virtual-navigation decision task. Nature 484, 62–68 (2012). 10.1038/nature10918

[7] Keller, G. B., Bonhoeffer, T. & Hubener, M. Sensorimotor mismatch signals in primary visual cortex of the behaving mouse. Neuron 74, 809–815 (2012). 10.1016/j.neuron.2012.03.040

[8] Lopes, G. et al. BonVision–an open-source software to create and control visual environments. BioRxiv, 2020.2003. 2009.983775 (2020).

[9] Stringer, C. et al. Spontaneous behaviors drive multidimensional, brainwide activity. Science 364, eaav7893 (2019).

[10] Reimer, J. et al. Pupil fluctuations track fast switching of cortical states during quiet wakefulness. Neuron 84, 355–362 (2014). 10.1016/j.neuron.2014.09.033

[11] Vinck M, Batista-Brito R, Knoblich U, Cardin JA. Arousal and locomotion make distinct contributions to cortical activity patterns and visual encoding. Neuron. 2015;86(3):740–54.

[12] McGinley, M. J. et al. Waking state: rapid variations modulate neural and behavioral responses. Neuron 87, 1143–1161 (2015).

[13] Naik, H., Bastien, R., Navab, N. & Couzin, I. D. Animals in virtual environments. IEEE Transactions on Visualization and Computer Graphics 26, 2073–2083 (2020).

[14] Shimaoka, D. in Awake Behaving Mesoscopic Brain Imaging 185–208 (Springer, 2024).

[15] MacDowell, C. J. & Buschman, T. J. Low-Dimensional Spatiotemporal Dynamics Underlie Cortex-wide Neural Activity. Curr Biol 30, 2665–2680 e2668 (2020). 10.1016/j.cub.2020.04.090

[16] Steinmetz, N. A., Zatka-Haas, P., Carandini, M. & Harris, K. D. Distributed coding of choice, action and engagement across the mouse brain. Nature 576, 266–273 (2019). 10.1038/s41586-019-1787-x

[17] Ma, Y. et al. Wide-field optical mapping of neural activity and brain haemodynamics: considerations and novel approaches. Philosophical Transactions of the Royal Society B: Biological Sciences 371 (2016).

[18] Vanni, M. P. & Murphy, T. H. Mesoscale transcranial spontaneous activity mapping in GCaMP3 transgenic mice reveals extensive reciprocal connections between areas of somatomotor cortex. Journal of Neuroscience 34, 15931–15946 (2014).

[19] Grosenick, L., Marshel, J. H. & Deisseroth, K. Closed-loop and activity-guided optogenetic control. Neuron 86, 106–139 (2015). 10.1016/j.neuron.2015.03.034

[20] Zerbi, V. et al. Resting-state functional connectivity changes in aging apoE4 and apoE-KO mice. Journal of Neuroscience 34, 13963–13975 (2014).

[21] Carroll, L. et al. Autism spectrum disorders: multiple routes to, and multiple consequences of, abnormal synaptic function and connectivity. The Neuroscientist 27, 10–29 (2021).

[22] Nakai, N. et al. Virtual reality-based real-time imaging reveals abnormal cortical dynamics during behavioral transitions in a mouse model of autism. Cell Reports 42 (2023).

[23] Venkataramani, V. et al. Glioblastoma hijacks neuronal mechanisms for brain invasion. Cell 185, 2899–2917 e2831 (2022). 10.1016/j.cell.2022.06.054

